# Reprogramming of Lipid Metabolism by FLASH Radiotherapy Selectively Protects Radiosensitive Normal Tissues

**DOI:** 10.64898/2026.04.20.719755

**Authors:** Shuhei Aramaki, Maxime Dubail, Mathieu Sertorio, Md Muedur Rahman, Mst Sayela Afroz, Chi Zhang, Liang Tang, Kohei Wakabayashi, Kenta Konishi, Tomoaki Kahyo, Gael Boivin, Marta Vilalta, Ricky A Sharma, Katsumasa Nakamura, Mitsutoshi Setou

## Abstract

FLASH radiotherapy, delivered at ultra-high dose rates exceeding 100 Gy/s, spares normal tissues while maintaining tumor control, yet the molecular mechanisms underlying this differential response remain poorly understood. Here we employed spatial and bulk multi-omics to investigate lipid and protein remodeling in tongue tissue and Mouse oral carcinoma 2 (MOC2) tumors at two weeks post-irradiation with FLASH or conventional dose-rate (CONV) proton radiotherapy. Bulk lipidomics revealed that CONV irradiation induced marked triglyceride (TG) depletion in tongue tissue, whereas FLASH attenuated this depletion. Spatial lipidomics using MALDI-MSI demonstrated that this TG loss was spatially restricted to minor salivary glands, identifying these radiosensitive structures as focal points of radiation-induced lipid damage. Additionally, FLASH irradiation uniquely promoted increases in membrane phospholipids and their lysophospholipid intermediates, consistent with active phospholipid turnover rather than passive damage avoidance. Proteomics revealed divergent metabolic programs: CONV activated a destructive cascade characterized by Ces1d depletion, Acox1-mediated peroxisomal β-oxidation, and Acot7-driven fatty acid overflow, collectively defining a lipid droplet collapse; whereas FLASH engaged a protective program featuring Mgll suppression, Apoe-mediated triglyceride redistribution, and Lypla2-mediated lysophospholipid clearance. We propose a lipid metabolic reprogramming hypothesis in which FLASH not only minimizes acute oxidative damage but actively reprograms lipid metabolism to preserve lipid droplet stores and promote membrane remodeling. These findings provide a putative molecular foundation for the FLASH effect and support published data on the protection of normal tissues.

## Introduction

Radiotherapy is a central component of the management of head and neck cancers (HNC), but its clinical efficacy is limited by dose-dependent toxicity to surrounding normal tissues^1^. Irradiation of the oral cavity, salivary gland, and tongue frequently induces severe acute and chronic side effects, including oral mucositis, hypogeusia, xerostomia and fibrosis, which markedly reduce patients’ quality of life and may compromise treatment compliance^2–4^.

Ultra-high dose-rate radiotherapy, known as FLASH radiotherapy, has emerged as a promising strategy to widen the therapeutic window of radiotherapy in HNC. Delivered at dose rates exceeding 100 Gy/s, FLASH irradiation has been shown in multiple preclinical models to significantly reduce normal tissue toxicity while preserving tumor control when compared to conventional dose-rate radiotherapy^5^. These protective effects have been reported across several organs, including brain^6–11^, lung^12–14^, skin^15–18^, gastrointestinal tract^19–22^, heart^23^ and oral tissues^24–26^, positioning FLASH as a potentially transformative approach for clinical radiotherapy.

In the context of head and neck irradiation, the normal tissue–sparing effects of FLASH radiotherapy have been demonstrated in veterinary trials in cats and dogs with oral tumors^25,26^. Building on these veterinary observations, murine studies using proton FLASH radiotherapy showed a significant reduction in radiation-induced salivary gland dysfunction and oral mucositis, together with preserved tumor control, in orthotopic tongue tumor models^24^. While these studies clearly demonstrated functional and histopathological protection of highly radiosensitive oral tissues, the underlying molecular mechanisms were not explored. On the tumor side, Li et al. demonstrated using transcriptomics and proteomics that FLASH irradiation reverses radioresistance in HNSCC through tumor immune microenvironment activation in a MOC1 model^27^. Similarly, Ni et al. showed that FLASH radiation reprograms lipid metabolism in tumor-associated macrophages, abrogating lipid oxidase expression to reverse immunosuppression^28^. Thus, while the molecular effects of FLASH on tumors are increasingly characterized, the basis of the normal tissue-sparing effect remains largely unexplored.

Among the mechanisms proposed to explain the normal tissue-sparing effect of FLASH, alterations in redox chemistry and oxidative stress have emerged as putative key components as well as their impact on stem cell niches^12,19,29^. Ionizing radiation induces the generation of reactive oxygen species, leading to oxidative damage of cellular macromolecules, including lipids^30,31^. In particular, lipid peroxidation of polyunsaturated fatty acids within membrane phospholipids is a key mediator of radiation-induced cellular injury and inflammatory signaling^30,32–34^. Beyond generic lipid peroxidation, previous lipidomic work revealed an ultra-early remodeling of polyunsaturated fatty acid (PUFA) containing lipids in kidney, occurring within minutes after high-dose irradiation^34^. These observations support the concept that lipids can actively modulate radiation responses by transiently reshaping membrane composition, buffering reactive oxygen species, and initiating lipid-derived signaling cascades that influence inflammation and cell fate decisions, rather than acting as passive substrates of oxidative damage.

Recent studies point to lipid chemistry as a key determinant of the FLASH response at the molecular level. Using simplified membrane liposomal or micelle structure, Froidevaux et al. demonstrated that FLASH irradiation markedly suppresses lipid peroxidation compared with conventional irradiation^35^. Extending these findings to biologically active lipid mediators, Portier et al. showed that FLASH irradiation selectively down-regulates oxylipin production in normal cells and mouse lung tissue within minutes after exposure, an effect absent in tumor cells and mitigated by hypoxia, highlighting the importance of membrane composition and oxygen-dependent lipid signaling^36^. Finally, these lipid effects were linked to ferroptosis, demonstrating that FLASH irradiation limited lipid peroxidation and ferroptotic cell death in normal tissues while preserving ferroptosis induction in tumors, and that increasing iron availability in normal tissues abolished FLASH protection. Together, these studies suggest an important role of lipid composition and oxidation at acute timepoint post-treatment. However, lipid alterations occurring in the inflammatory and regenerative phase of tissue have not been explored.

We therefore investigated the molecular consequences of a single 10 Gy fraction delivered with proton FLASH (100 Gy/s) or conventional dose rate protons (1 Gy/s), focusing on the tongue as a clinically relevant normal tissue and on contralateral MOC2 tumors in a mouse model. We selected a two-week post-irradiation timepoint to interrogate molecular remodeling occurring beyond the immediate phase of radiation-induced damage, at a stage where inflammatory responses, tissue repair, and regenerative processes are actively engaged. We integrated bulk lipidomic and proteomic profiling using LC–MS/MS with mass spectrometry imaging (MSI) to capture both the molecular and spatial dimensions of radiation-induced remodeling. This multimodal approach enabled the characterization of coordinated lipid and protein alterations, while resolving their tissue- and structure-specific distributions. By jointly interrogating tumor and normal tissues at this post-acute stage, we aimed to identify lipid- and protein-driven pathways that may underlie the differential tissue responses to FLASH versus conventional radiotherapy.

## Material and methods

### Cell culture

MOC2 mouse oral carcinoma cells (Kerafast) were detached with TrypLE Express (Gibco) for 3 min at 37 °C. Cells were resuspended in IMDM-based MOC medium containing 62.6% Iscove’s Modified Dulbecco’s Medium (IMDM) 1X (Gibco), 31.3% Ham’s F-12 Nutrient Mixture 1X (Gibco), 5% fetal bovine serum (Corning), 1% penicillin/streptomycin (Gibco), 5 mg/mL insulin (Sigma-Aldrich), 0.04 μg/mL hydrocortisone (Sigma-Aldrich), and 5 ng/mL EGF (Merck Millipore). The medium was sterilized with a 1 L vacuum filter/storage bottle system fitted with a 0.22 µm PES membrane (Corning). Cells were maintained in a humidified incubator at 37 °C with 5% CO2. Experiments were performed when cultures reached 80% confluence. Cultures were tested routinely for mycoplasma by PCR.

### Ethical consent and animal experiments

All animal experiments were approved by the Cincinnati Children’s Hospital Institutional Animal Care and Use Committee (IACUC2023-0013). All procedures were conducted in compliance with the National Institutes of Health (NIH) Guide for the Care and Use of Laboratory Animals and followed the ARRIVE (Animal Research: Reporting of In Vivo Experiments) guidelines.

Ten-week-old female C57BL/6 mice (The Jackson Laboratory, Bar Harbor, ME, USA) were acclimated for 7 days before experimentation and housed in the animal facility of Cincinnati Children’s Medical Center. Mice were housed in groups of 4 per cage under controlled conditions (21.1 ± 1 °C, 50 ± 5% humidity, and a 12-hour light/dark cycle) with free access to food and tap water.

### Orthotopic tumor model

Orthotopic tumor injection was performed as described previously^37^. Briefly, anesthetized mice received a 100 µL injection of MOC2 cell suspension into the right buccal epithelium. The suspension contained 0.5 × 10[ MOC2 cells in 50 µL PBS mixed with 50 µL Matrigel (354234, Corning). Anesthesia was induced with 4% isoflurane and maintained at 2.5% using a SomnoSuite system (Kent Scientific) with a nose cone (Kent Scientific), which allowed access to the right oral commissure. Tumors were allowed to grow for 28 days before irradiation. Mice were then randomized into three groups: control (sham-irradiated), conventional dose-rate irradiation (CONV), and ultra-high dose-rate irradiation (FLASH). Both tumor-bearing and non-tumor-bearing mice received a single fraction of 10 Gy to the head and neck region as described in the Proton Delivery section. Sample sizes were n = 3 per group.

### Tissue collection

Two weeks (14 days) after irradiation, mice were euthanized in their housing cage by CO2 inhalation using an automatic air displacement container connected to a CO2 compressed gas cylinder. Mice were maintained until breathing stopped, followed by at least 5 additional minutes of exposure, followed by cervical dislocation as a secondary method to ensure euthanasia. Tongue tissues and tumors were harvested and immediately snap-frozen in liquid nitrogen for bulk lipidomics and proteomics analyses, or embedded in Super Cryoembedding Medium (SCEM; SECTION-LAB Co., Ltd., Hiroshima, Japan) for cryosectioning and spatial lipidomics analysis. Samples were stored at −80 °C until further processing.

### Proton Delivery, Dosimetry and Monitoring

The ProBeam Pencil Beam Scanning Gantry (Varian Medical Systems, Palo Alto, CA, USA) was used to deliver a monoenergetic, single-layer transmission radiation field. FLASH dose rates were delivered at 250 MeV, while conventional dose rates were delivered at 244 MeV. Since the water-equivalent depth of the mouse is small (i.e., approximately 1.0 cm), the small change in incident energy does not yield a measurable difference in linear energy transfer or material stopping power at the mouse depth. Conventional irradiation was delivered at a dose rate of 1 Gy/s, while FLASH irradiation was delivered at a dose rate of 100 Gy/s.

The field dosimetric metrology system is composed of both an ion chamber and electrometer and radiochromic film and scanner. Ion chamber and electrometer calibration factors have been determined independently by an Accredited Dosimetry Calibration Laboratory (ADCL). The radiochromic film and scanner are cross-calibrated to the ion chamber. Using the International Atomic Energy Agency (IAEA) TRS-398 absolute dose formalism, collection efficiency and recombination effects were quantified using two-voltage technique, and the ion chamber correction was measured to be less than 1.0% in all conditions. Independently, the ion chamber was validated against a graphite calorimeter in order to ensure accuracy of dose and dose-rate independence. The ion chamber and graphite calorimeter agree within 1.0%. Fields were measured for flatness and symmetry using calibrated radiochromic film (Gafchromic EBT3; Ashland, Bridgewater, NJ, USA) and calibrated flatbed scanner (Epson 10000XL). Furthermore, fields were measured for absolute dose and dose rate using a calibrated Advanced Markus ion chamber (PTW, Freiburg im Breisgau, Germany) connected to a calibrated IBA Dose1 electrometer (IBA-Dosimetry, Schwarzenbruck, Germany). Tolerances of 5% flatness and symmetry, 3% for dose and 5% for dose rate were used.

The single-layer spot patterns were designed using in-house software to deliver a uniform dose of 10 Gy (within 5% uniformity) to a field of 25 mm × 23 mm (95% isodose line) composed of 30 individual spots (beam spot size σ = 4.0 mm) at a 0 degree gantry position. The field was irradiated with a continuous beam without beam pause between the spots. Prior to irradiation, the dose was measured at a water-equivalent depth of 1.0 cm in a solid water phantom using a calibrated Advanced Markus ion chamber connected to a calibrated IBA Dose1 electrometer. Dose rate was determined by computing the ratio of the total dose to total field delivery time as provided by the machine log files. During irradiation, an Advanced Markus ion chamber was placed distal to the mouse to verify the dose for each irradiation.

To ensure proper alignment, the mouse jig was first positioned at isocenter using the gantry laser alignment system (accuracy 1 mm). The radiation field alignment was visually verified by irradiating radiochromic film secured within the jig. Each mouse was placed such that the tongue was coinciding with the isocenter using both the laser system and visual indication from the radiochromic film. Localization was verified on a subset of mice using the orthogonal kilovoltage image guidance system, and no discrepancies with the laser alignment were detected. A summary of all irradiation parameters is provided in Supplementary Table 1.

### Lipid extraction and LC-MS/MS measurements

Lipid extraction from mouse tongue tissue was performed using a modified Bligh and Dyer protocol^38^. Tissue slices were processed using a solvent ratio of 2:2:1.8 (methanol:chloroform:water). Briefly, samples were first homogenized by adding an appropriate volume of ice-cold methanol to ensure comparable sample concentrations, followed by vortexing and sonication using a BioRaptor system (Bio-Rad, USA). Chloroform was then added in two sequential steps. In the first step, half of the total chloroform volume was added, vortexed thoroughly, and incubated for 10–15 min at room temperature. Subsequently, the remaining chloroform volume containing the internal standard PC(12:0/12:0) (10 ng/µL) was added and mixed thoroughly by vortexing. Phase separation was induced by adding 0.28 M acetate solution according to the water ratio, followed by vortexing and centrifugation at 4,000 rpm for 30 min at 4 °C. The lower organic phase was carefully collected, dried using a vacuum concentrator, and reconstituted in 50–100 µL of 100% methanol prior to LC–MS/MS analysis. Lipidomic analyses were performed using a Q Exactive Orbitrap mass spectrometer (Thermo Fisher Scientific, USA) coupled to a liquid chromatography system. Chromatographic separation was achieved using an Acclaim 120 column (150 mm × 2.1 mm, 3 µm; Thermo Fisher Scientific). The mobile phases consisted of solvent A (water:acetonitrile:methanol, 2:1:1, v/v/v) and solvent B (acetonitrile:isopropanol, 1:9, v/v), both supplemented with 5 mM ammonium formate and 0.1% formic acid. The gradient was programmed as follows: 20% B at 0 min, increased linearly to 100% B over 50 min, held at 100% B for 10 min, then returned to 20% B over 10 min, for a total run time of 70 min. The flow rate was set to 0.30 mL/min, and the injection volume was 1 µL. Electrospray ionization (ESI) was performed in both positive and negative ion modes. The column temperature was maintained at 50 °C. Source parameters were set as follows: spray voltage 3.5 kV, capillary temperature 250 °C, heater temperature 350 °C, sheath gas flow rate 50, auxiliary gas flow rate 15, and S-lens RF level 50. Mass spectra were acquired over an m/z range of 220–2000 at a resolution of 70,000.

### Protein extraction and LC-MS/MS measurements

Proteins were extracted from mouse tongue tissue slices using a chaotropic lysis buffer containing 6 M urea, 2 M thiourea, 0.4% SDS, and 1 mM dithiothreitol (DTT). Protein precipitation was performed by adding ice-cold acetone at a 1:4 (sample:acetone, v/v) ratio, followed by overnight incubation at −20 °C. Samples were then centrifuged at 15,000 rpm for 10 min to collect the protein pellet.Protein pellets were resuspended in 0.5 M triethylammonium bicarbonate (TEAB; Sigma-Aldrich) buffer using gentle sonication. Protein concentration was determined using a Bradford assay (Bradford Reagent Plus, Thermo Fisher Scientific) according to the manufacturer’s instructions. For each sample, 10 µg of protein was aliquoted into low-binding tubes, and the volume was adjusted to 100 µL with 50 mM ammonium bicarbonate buffer. Proteins were reduced with 10 mM DTT (final concentration) at 95 °C for 15 min, followed by alkylation with 50 mM iodoacetamide (final concentration; Thermo Fisher Scientific) at room temperature in the dark for 30 min. Samples were subsequently diluted with 50 mM ammonium bicarbonate to reduce the TEAB concentration to below 50 mM prior to enzymatic digestion. Proteolytic digestion was performed sequentially using LysC and trypsin. Samples were first incubated with LysC (MS grade, Promega) at a 1:100 (enzyme:protein, w/w) ratio at 37 °C for 2–3 h. Trypsin (MS grade, Promega) was then added at a 1:50 (enzyme:protein, w/w) ratio and digestion was continued at 37 °C for 10 h with gentle agitation. Digestion was terminated by acidification with 1% trifluoroacetic acid (TFA; Sigma-Aldrich). Peptides were desalted using MonoSpin C18 columns (GL Sciences) according to the manufacturer’s protocol and dried using a vacuum concentrator. Dried peptides were reconstituted in 0.1% formic acid prior to LC–MS/MS analysis. Peptide separation and analysis were performed using a Q Exactive Orbitrap mass spectrometer (Thermo Fisher Scientific) coupled to a nano-liquid chromatography system. Peptides were separated on an NTCC-360/75-3-125 column (Nikkyo Technos Co., Japan) using a binary solvent system consisting of solvent A (0.1% formic acid in water) and solvent B (80% acetonitrile in 0.1% formic acid). The LC gradient was programmed as follows: 0–35% B over 50 min, 35–100% B over 5 min, followed by a 10 min hold at 100% B, resulting in a total run time of 65 min. The flow rate was set to 300 nL/min, and the injection volume was 1 µL. Electrospray ionization was performed in positive ion mode with the following parameters: spray voltage 4.3 kV, capillary temperature 320 °C, sheath gas flow rate 30, auxiliary gas flow rate 5. Data were acquired in both data-dependent acquisition (DDA) and data-independent acquisition (DIA) modes. Full MS scans were acquired at a resolution of 70,000, with a scan range of m/z 350–1800 for DDA and m/z 350–980 for DIA analyses.

### Lipidomic MALDI-MSI measurements

Serial cryosections of mouse tongues (10 µm thickness) were prepared using a cryomicrotome (Leica CM1950, Germany) and mounted onto indium tin oxide (ITO)– coated glass slides (Matsunomi, Japan). Tissue sections were dried under vacuum in a desiccator for 30 min prior to matrix application. Matrix deposition was performed using an automated TM sprayer (HTX Technologies, USA), with separate matrices applied for positive and negative ion mode acquisitions. For positive ion mode, 2,5-dihydroxybenzoic acid (DHB; Sigma-Aldrich) was prepared at 40 mg/mL in methanol:water (1:1, v/v) and sprayed onto tissue sections under the following conditions: matrix flow rate 0.10 mL/min, nozzle temperature 80 °C, gas pressure 10 psi, gas flow rate 3 L/min, nozzle velocity 1250 mm/min, 8 passes, and criss-cross spray pattern. For negative ion mode acquisition, 9-aminoacridine (9-AA; Sigma-Aldrich) was prepared at 10 mg/mL in ethanol:water (7:1, v/v) and applied using the following parameters: matrix flow rate 0.12 mL/min, nozzle temperature 90 °C, gas pressure 10 psi, gas flow rate 3 L/min, nozzle velocity 1250 mm/min, 7 passes, and criss-cross spray pattern. MALDI–MSI data were acquired using a Fourier transform ion cyclotron resonance (FT-ICR) mass spectrometer (Solarix 7T, Bruker Daltonics, Germany) equipped with both electrospray ionization (ESI) and MALDI sources. Prior to data acquisition, external calibration was performed using sodium formate (1 mg/mL in methanol:water, 95:5, v/v) via the ESI source. MSI acquisition parameters were as follows: mass range m/z 100–1200, positive and negative ion polarities, data size 1M, scan rate 1 scan/s, laser power 23%, 100 laser shots per pixel, laser frequency 1000 Hz, small laser diameter, and raster step size of 50 µm.

### Peptides MALDI-MSI measurements

Tissue sections were prepared for peptide mass spectrometry imaging (MSI) following on-tissue tryptic digestion. Briefly, mounted tissue sections were first delipidated by sequential washes in 100% ethanol (twice), 80% ethanol, 70% ethanol, and ultra-pure water, each for 2 min and repeated twice. Slides were then dried under vacuum in a desiccator for 5 min. Antigen retrieval was performed by autoclaving the slides at 100 °C for 30 min in 10 mM ammonium citrate buffer, followed by gentle buffer exchange with ultra-pure water. On-tissue enzymatic digestion was carried out by spraying MS-grade trypsin (Promega, USA) using a pneumatic TM sprayer (HTX Technologies). Trypsin was prepared at a concentration of 0.074 mg/mL in 100 mM ammonium bicarbonate containing 9% acetonitrile and applied under the following conditions: flow rate 8 µL/min, nozzle temperature 30 °C, gas pressure 10 psi, gas flow rate 3 L/min, nozzle velocity 750 mm/min, track spacing 1.5 mm, 8 spray cycles, and criss-cross spray pattern. One tissue section was covered with aluminum foil and processed in parallel as a non-digested control. Slides were incubated in a humid chamber at 37 °C for 2.5 h to allow peptide generation. Following digestion, slides were dried again under vacuum for 30 min prior to matrix application. α-Cyano-4-hydroxycinnamic acid (CHCA; Sigma-Aldrich) was prepared at 5 mg/mL in acetonitrile:water (1:1, v/v) containing 0.1% trifluoroacetic acid and deposited using the TM sprayer with the following parameters: matrix flow rate 0.10 mL/min, nozzle temperature 80 °C, gas pressure 10 psi, gas flow rate 3 L/min, nozzle velocity 1250 mm/min, and 8 spray cycles in a criss-cross pattern. Peptide MSI data were acquired using a Solarix 7T Fourier transform ion cyclotron resonance (FT-ICR) mass spectrometer (Bruker Daltonics, Germany). Prior to acquisition, the instrument was externally calibrated using sodium formate (1 mg/mL in methanol:water, 95:5, v/v). Data were acquired in positive ion mode over an m/z range of 500–3000, with a data size of 512k, laser power set to 40%, 100 laser shots per pixel, a laser frequency of 1000 Hz, small laser focus, and a raster step size of 50 µm.

### Data analysis and statistical analysis

LC-MS/MS for lipidomic data was analyzed by Xcaliber^TM^ (Thermo Fisher) and LipidSearch^TM^ (Thermo Fisher). Lipidomic data was normalized by internal standard [PC 12:0/12:0]. Raw lipidomic data were processed in R (v.4.2) using a standardized pipeline. Lipid species with excessive missing values were excluded using a strict filter (more than one missing value per condition). When multiple features corresponded to the same lipid annotation, only the most intense feature was retained. Intensities were first normalized on the linear scale using the internal standard PC(12:0/12:0) to correct for extraction and analytical variability. Internal standard–normalized values were then normalized using a median-of-ratios (MoR) approach, in which sample-specific size factors were calculated from the median ratio of lipid intensities to lipid-wise geometric means computed across a reference set detected in at least 50% of samples. Normalized data were log2-transformed and remaining missing values were imputed using a minimum detection (MinDet) approach. A final biochemical filtering step was applied to retain mammalian-relevant lipid species based on lipid class, acyl chain number, chain length, and degree of unsaturation. The resulting dataset was used for all downstream analyses. Differential analysis was performed on IS + MoR–normalized log[ lipid intensities using linear models with empirical Bayes moderation (limma). Comparisons were conducted per tissue between CONV vs CTRL, FLASH vs CTRL, and FLASH vs CONV conditions. Lipids with *p* < 0.05 and |log[FC| > 0.5 were considered significantly regulated. For the analysis of bottom-up proteomic data in DIA acquisition, we used DIA-NN software^39^. Quantitative proteomic data were processed using a workflow conceptually similar to the lipidomic analysis pipeline, including quality filtering, normalization, log[ transformation, and differential analysis, but without the use of an internal standard. MSI data were acquired by ftms control^TM^ (Bruker Daltonics) and fleximaging^TM^ (Bruker Daltonics). Later, the MSI data were exported as imZML file and analyzed by Imagereveal^TM^ software (Shimadzu, Japan).

### Annotation of MALDI-MSI data

Accurate-mass lipid/peptide features detected by MALDI-FT-ICR-MSI (Solarix 7T) were annotated by matching experimental *m/z* values to a curated lipid reference list derived from LC–MS/MS measurements. For each analyte, theoretical *m/z* values were generated in silico for the main expected adducts in positive and negative ion modes. MSI *m/z* values from each sheet were matched to the adducted theoretical masses within a tolerance of ±10 ppm, and all candidates within this window were retained and ranked by absolute mass error (Δppm). To minimize matrix/background-driven features, a matrix-only reference file acquired under the same conditions was used for feature filtering. Reproducible MSI features were then defined across three technical replicates per polarity.

### Histological staining

Following MALDI-MSI acquisition, matrix was removed from the tissue sections by immersing slides in 80% acetone for 10 seconds without agitation, followed by a second 80% acetone bath for 10 seconds. Slides were briefly rinsed with ultrapure water and then fixed in 4% paraformaldehyde (PFA) for 15 minutes. After washing under running tap water for approximately 10 minutes, sections were stained with Mayer’s hematoxylin for 5 minutes, washed under running tap water for 5 minutes, and then placed in warm water (approximately 40°C) for 5 minutes for bluing. Sections were subsequently stained with ethanol-based eosin for 3 minutes. Excess stain was rinsed off with tap water, and slides were dehydrated through a graded ethanol series (80%, 100%, 100%, 100%), cleared in three changes of xylene, and coverslipped with a non-aqueous mounting medium. Hematoxylin and eosin staining times were optimized according to institutional standard protocols. H&E-stained sections were used for histological annotation and spatial co-registration with MALDI-MSI lipid images.

## Results

### CONV versus FLASH irradiation reveals strong lipidomic divergence in irradiated tongue

MOC2 tumor–bearing mice received a single 10 Gy fraction delivered at either conventional (CONV) or ultra-high dose-rate FLASH protons, with non-irradiated animals used as controls (CTRL) (Fig. 1A). Tongue tissue and tumors were collected two weeks post-irradiation and analyzed using complementary bulk lipidomics and spatial lipidomic approaches (Fig. 1A). After standardized data processing and biochemical filtering to retain mammalian-relevant lipid species (Supplementary Fig. 1A), a total of 1,547 lipid species spanning 37 lipid classes were quantified across tissues and experimental conditions (Supplementary Fig. 1B–C). Analysis of lipid unsaturation patterns further revealed class-specific distributions of saturated, monounsaturated, and polyunsaturated fatty acyl chains, highlighting the biochemical diversity of the detected lipid species and supporting the coverage of structurally and functionally distinct lipid subclasses (Supplementary Fig. 1C).

**Figure 1:**
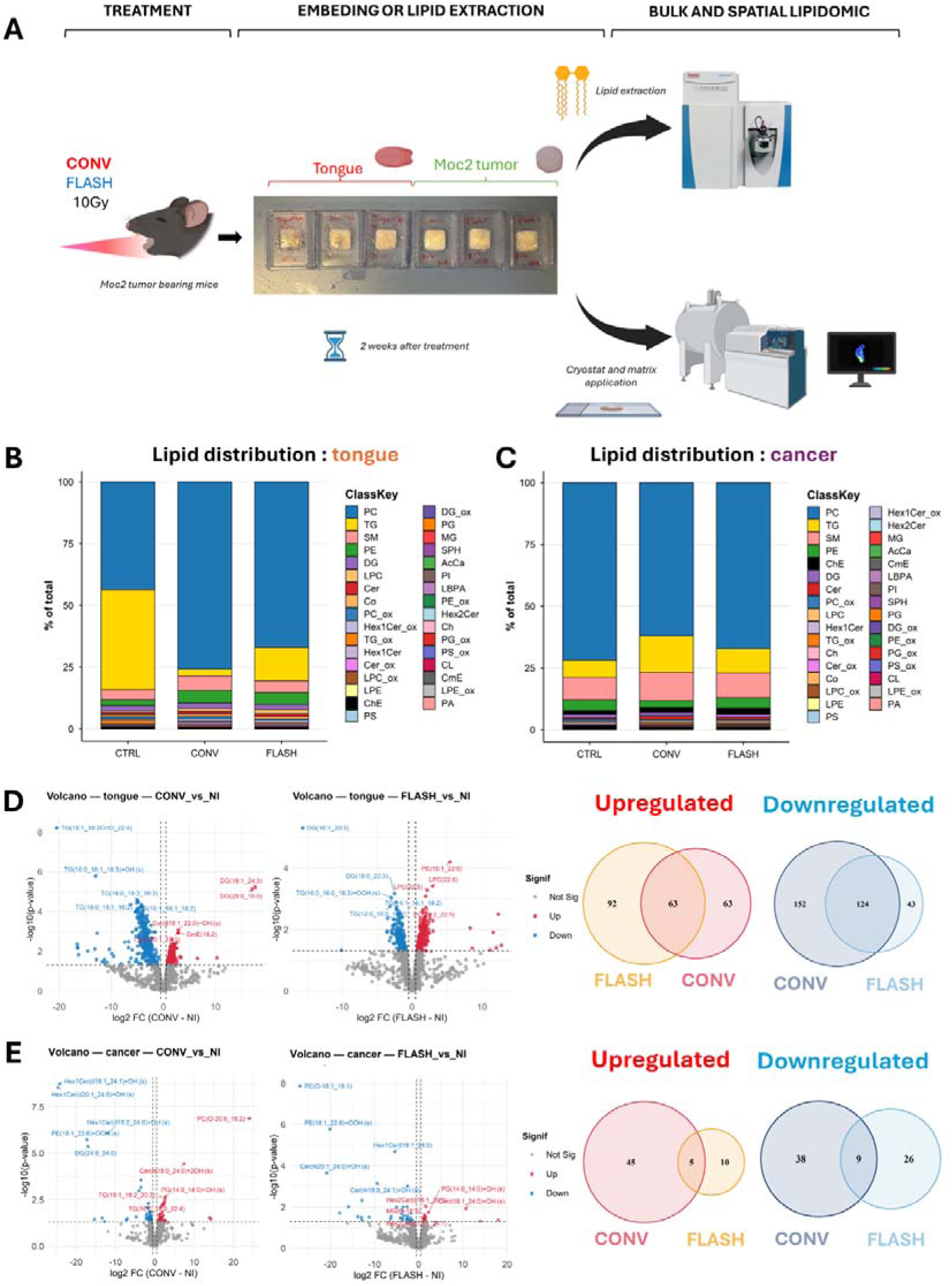
LC-MS/MS lipidomic profiling reveals tissue-specific responses after conventional and FLASH irradiation. (A) Scheme of experimental design. MOC2 tumor-bearing or non-tumor-bearing mice received a single 10 Gy irradiation delivered at either conventional dose rate (CONV) or ultra-high dose-rate FLASH. Non-irradiated animals served as controls (CTRL). Tongue tissue and contralateral MOC2 tumors were harvested two weeks post-irradiation for bulk lipidomic analysis by LC-MS/MS and spatial lipidomic analysis by mass spectrometry imaging (MSI). (B) Lipid class distribution in tongue tissue across experimental conditions. Stacked bar plots show the relative abundance (% of total) of major lipid classes in CTRL, CONV, and FLASH groups. (C) Lipid class distribution in MOC2 tumor tissue. Stacked bar plots show limited changes in lipid class composition across experimental conditions. (D) Differential lipid analysis in tongue tissue. Left: Volcano plots showing significantly regulated lipid species comparing CONV vs CTRL and FLASH vs CTRL conditions. Right: Venn diagrams displaying the number of upregulated and downregulated lipid species. (E) Differential lipid analysis in tumor tissue. Left: Volcano plots showing limited differential regulation in tumors. Right: Venn diagrams displaying the number of upregulated and downregulated lipid species.

Baseline lipid class distributions revealed marked tissue-specific differences (Fig. 1B–C). Tongue tissue exhibited a higher relative contribution of neutral lipids, particularly triglycerides that represent 40.5% against 14.8 % in tumor (TG). Tumors were enriched in membrane-associated lipid classes, including phosphatidylcholines representing 71.9% against 43.6 % in tongue (PC) and sphingolipids with 9.1 % against 4.0%. Upon irradiation, tongue tissue underwent pronounced lipidomic remodeling, marked by a substantial reduction in triglyceride (TG) abundance. This decrease was markedly stronger following CONV irradiation (− 37.7% vs CTRL) than after FLASH (− 27.1% vs CTRL) (Fig. 1B). In contrast, tumor tissues displayed only modest changes in lipid class composition following irradiation, with a limited reduction in phosphatidylcholine (PC) abundance after both CONV (− 9.8% vs CTRL) and FLASH (− 4.64% vs CTRL) irradiation, and minimal global lipid redistribution across conditions (Fig. 1C). Multivariate analysis performed within each tissue further supported these observations. In tongue tissue, lipidomic profiles separated according to irradiation condition, whereas tumors showed limited discrimination between CTRL, CONV, and FLASH samples (Supplementary Fig. 1D–E).

At the species level, differential analysis confirmed a substantially higher number of significantly regulated lipid species in tongue (537 species) tissue compared to tumors (133 species) (Fig. 1D–E). CONV irradiation affected a larger number of lipid species overall than FLASH irradiation, encompassing both up- and down-regulated species (CONV: up [n = 126], down [n = 276]; FLASH: up [n = 155], down [n = 167]). In tongue tissue, a subset of lipid species was commonly regulated by both irradiation modalities (up [n = 63], down [n = 124]), alongside modality-specific alterations (CONV-specific: up [n = 63], down [n = 152]; FLASH-specific: up [n = 92], down [n = 43]), indicating partially overlapping but quantitatively distinct lipid remodeling programs. In contrast, tumors exhibited markedly fewer significantly regulated lipid species (up [n = 66], down [n = 73]), with most changes being condition-specific (CONV-specific: up [n = 45], down [n = 38]; FLASH-specific: up [n = 10], down [n = 26]) and lacking coordinated patterns.

### The marked decrease of triglycerides in healthy tongue tissue following CONV is only partially observed after FLASH

Given the limited number of significantly regulated lipid species observed in tumors (n = 133); Supplementary Fig. 2A), subsequent analyses focused on healthy tongue tissue, where CONV and FLASH irradiation induced pronounced and divergent lipidomic responses (Fig. 2A). Among lipid classes showing increased abundance after irradiation, the classes were limited in number and predominantly involved membrane-associated lipid classes. Phosphatidylcholines (PC; n = 49), phosphatidylethanolamines (PE; n = 52), lysophosphatidylcholines (LPC; n = 15), lysophosphatidylethanolamine (LPE; n = 23) formed the core of this upregulated signature, consistent with selective remodeling of membrane lipid composition (Figure 2A). This pattern was further supported by functional enrichment analyses, which highlighted glycerophospholipids, zwitterionic headgroups, and intracellular membrane compartments and components, indicating coordinated membrane reorganization following irradiation (Figure 2B). Notably, this membrane-associated response was more prominently engaged after FLASH irradiation than after CONV conditions, with a substantially higher proportion of significantly upregulated membrane lipids (e.g., PC: 19.1% vs 14.2%; PE: 28.0% vs 10.9%; LPC: 42.1% vs 5.3%; LPE: 39.1% vs 13.0%) (Figure 2C).

**Figure 2:**
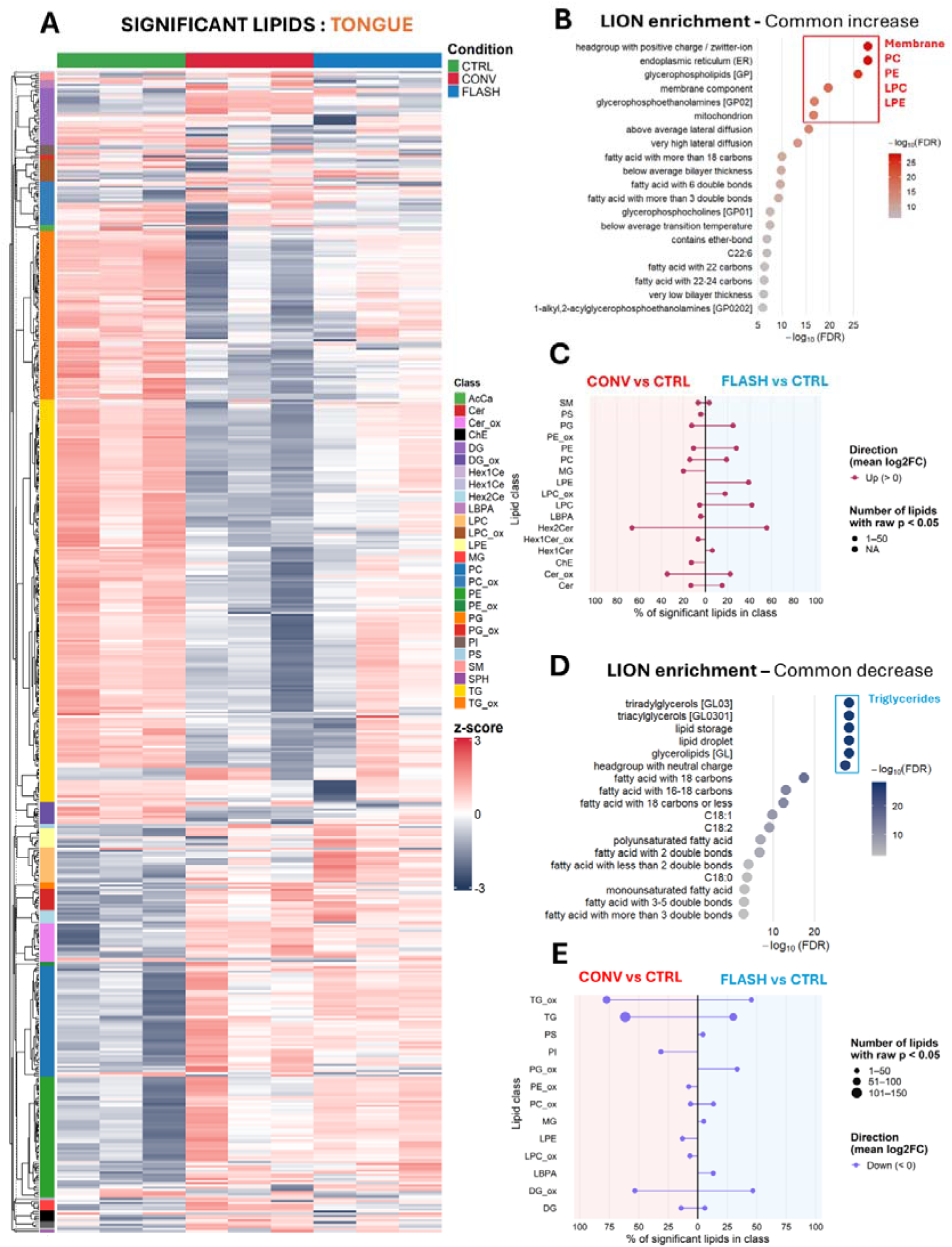
CONV irradiation induces coordinated triglyceride depletion and membrane lipid remodeling in tongue tissue. (A) Lipid class based hierarchical clustering heatmap of significantly regulated lipid species in tongue tissue across CTRL, CONV, and FLASH conditions. Lipids are annotated by class (color-coded sidebar). **(B)** LION (Lipid Ontology) enrichment analysis of commonly upregulated lipids after radiation. **(C)** Lipid class-level analysis of upregulated species. Butterfly plots show the percentage of significantly upregulated lipids within each class for CONV vs CTRL and FLASH vs CTRL comparisons. **(D)** LION enrichment analysis of commonly downregulated lipids after irradiation. **(E)** Lipid class-level analysis of downregulated species.

In contrast, the downregulated lipidomic response was broader and dominated by neutral lipid depletion. Triglycerides (TG; n = 180) and oxidized triglycerides (TG_ox; n = 78) accounted for most significantly altered species, together representing 81% of all downregulated lipids. (Figure 2A). Lipid ontology enrichment analyses revealed a strong overrepresentation of lipid storage–related features, including triglycerides, neutral lipids, lipid droplets, and energy reserve–associated terms, consistent with a coordinated loss of lipid storage compartments in tongue tissue (Figure 2D). Importantly, this depletion was substantially more pronounced after CONV irradiation, with 61.5% of total triglycerides (TG) and 77.3% of total oxidized triglycerides (TG_ox) significantly downregulated relative to CTRL. In contrast, FLASH irradiation induced a more limited reduction of these neutral lipid pools (28% TG and 45.4% TG_ox), indicating partial preservation of triglyceride stores after ultra-high dose-rate conditions.

By contrast, tumors showed a markedly attenuated lipidomic response to irradiation, with few lipid species significantly regulated after CONV or FLASH exposure and no coordinated class-level organization (Supplementary Figure 2A). Triglycerides were not depleted but instead displayed modest increases after both conditions, indicating a tongue-specific lipid response to radiation therapy.

To determine whether these bulk lipidomic alterations were spatially organized within the tissue, we next applied mass spectrometry imaging (MSI) at a 50-um resolution to resolve the regional distribution of radiation-responsive lipids in both tongue and tumor sections after FLASH and CONV dose rates.

### Spatial MSI reveals selective depletion of triglyceride species in minor salivary glands following CONV irradiation

To investigate whether the strong neutral lipid depletion observed by bulk LC-MS/MS was spatially organized within tongue tissue, we next examined the regional distribution of radiation-responsive triglyceride (TG) species using MALDI-MSI at 50-µm spatial resolution (Figure 3A). Among TG species significantly downregulated at the bulk level, only a subset exhibited sufficiently intense and spatially coherent ion images to allow confident MSI-based analysis (Figure 3B). These MSI-detectable species included multiple long-chain TGs, such as TG(16:1_18:1_18:2), TG(16:0_18:1_18:1), TG(18:1_18:2_18:2), and TG(18:1_18:1_18:2), which were representative of the broader TG class depleted after irradiation (Figure 3C).

**Figure 3.**
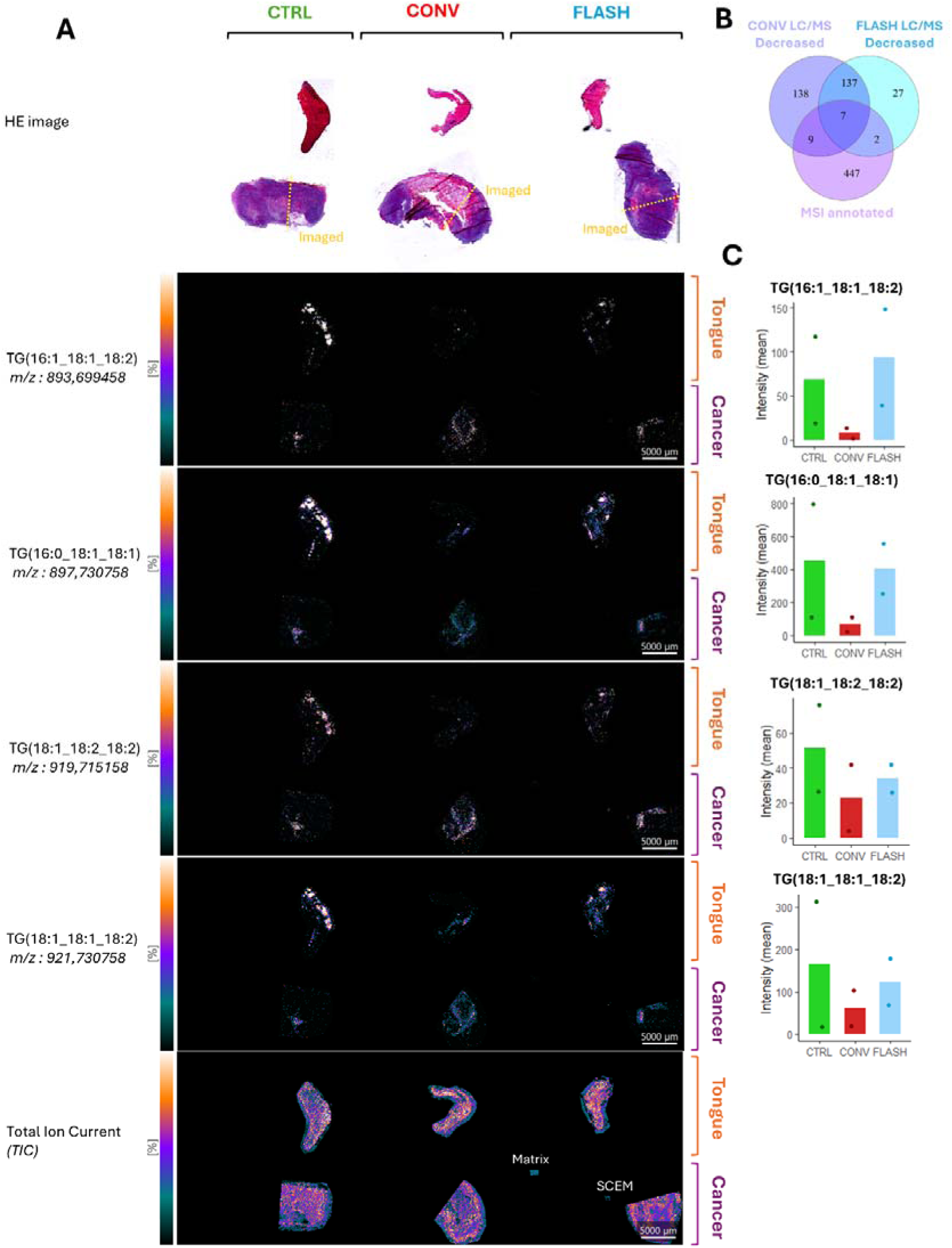
Spatial MSI reveals region-specific depletion of triglycerides in tongue tissue following irradiation, attenuated under FLASH conditions. (A) Upper : Representative H&E-stained serial sections of tongue tissue from CTRL, CONV-, and FLASH-irradiated mice, indicating the regions selected for MALDI-MSI acquisition (yellow dashed lines). Lower : MALDI-MSI ion images of representative downregulated TG species, including TG(16:1_18:1_18:2), TG(16:0_18:1_18:1), TG(18:1_18:2_18:2), and TG(18:1_18:1_18:2), displayed for tongue and tumor tissues under CTRL, CONV, and FLASH conditions. Ion intensity is shown using a normalized color scale. (B) Venn diagram showing the overlap between triglyceride (TG) species significantly decreased by bulk LC-MS/MS analysis after CONV or FLASH irradiation and those successfully annotated by MSI. (C) Quantification of mean signal intensity confirms a marked reduction of TG signals in tongue tissue following CONV irradiation, with partial preservation under FLASH irradiation.

Triglycerides (TG) are the major neutral lipid storage form in mammalian tissues and play a central role in buffering fatty acid availability, supporting membrane biosynthesis, and sustaining energy-demanding processes^40^. In the tongue, and particularly within minor salivary glands, TG-rich lipid droplets are abundant in serous and mucous acinar cells, reflecting the high secretory activity of these structures and their reliance on neutral lipid stores for membrane turnover and secretion-associated metabolism^41^.

Spatial ion maps revealed that TG signals in CTRL tongue tissue were not uniformly distributed but were strongly enriched within discrete glandular compartments. Overlay with H&E-stained sections demonstrated that these TG species were highly concentrated in minor salivary glands, encompassing both serous and mucous acini, consistent with the lipid-rich secretory nature of these structures (Figure 3A; Supplementary Figure 3). Following irradiation, a marked and spatially restricted loss of TG signal was observed within these glandular regions. This depletion was particularly pronounced after CONV irradiation, where TG signals were strongly reduced or nearly absent in minor salivary glands. In contrast, FLASH irradiation was associated with a partial preservation of TG signal intensity within the same acinar compartments, indicating attenuation of radiation-induced neutral lipid loss after ultra-high dose-rate conditions (Figure 3C).

Outside glandular regions, including epithelial and muscle compartments, TG signals were comparatively low at baseline and showed limited modulation across conditions. Together, these data identify minor salivary glands as a major anatomical locus of radiation-induced triglyceride depletion in tongue tissue and highlight their pronounced vulnerability to conventional irradiation, with partial protection preserved after FLASH conditions.

### Proteomic evidence for enhanced lipid catabolism after CONV irradiation compared with FLASH

To determine whether the contrasting lipidomic phenotypes induced by CONV and FLASH irradiation were underpinned by distinct protein-level programs, we next analyzed bulk proteomic profiles of tongue and cancer tissue (Figure 4). Differential analysis revealed a pattern distinct from that observed at the lipidomic level, with tumor tissue exhibiting a stronger proteomic response to irradiation than tongue tissue (Fig. 4A–B), dominated by wound healing, coagulation, and extracellular matrix organization pathways shared by both CONV and FLASH irradiation (Supplementary Figure 4A), indicating that tumor tissue did not exhibit modality-specific divergence. By contrast, in tongue tissue, CONV and FLASH irradiation each induced distinct proteomic responses relative to non-irradiated controls (Figure 4A). CONV irradiation differentially regulated 56 proteins (up [n = 31], down [n = 25]), whereas FLASH irradiation affected 52 proteins (up [n = 29], down [n = 23]), indicating a restricted yet structured set of radiation-responsive proteins in healthy tongue tissue.

**Figure 4.**
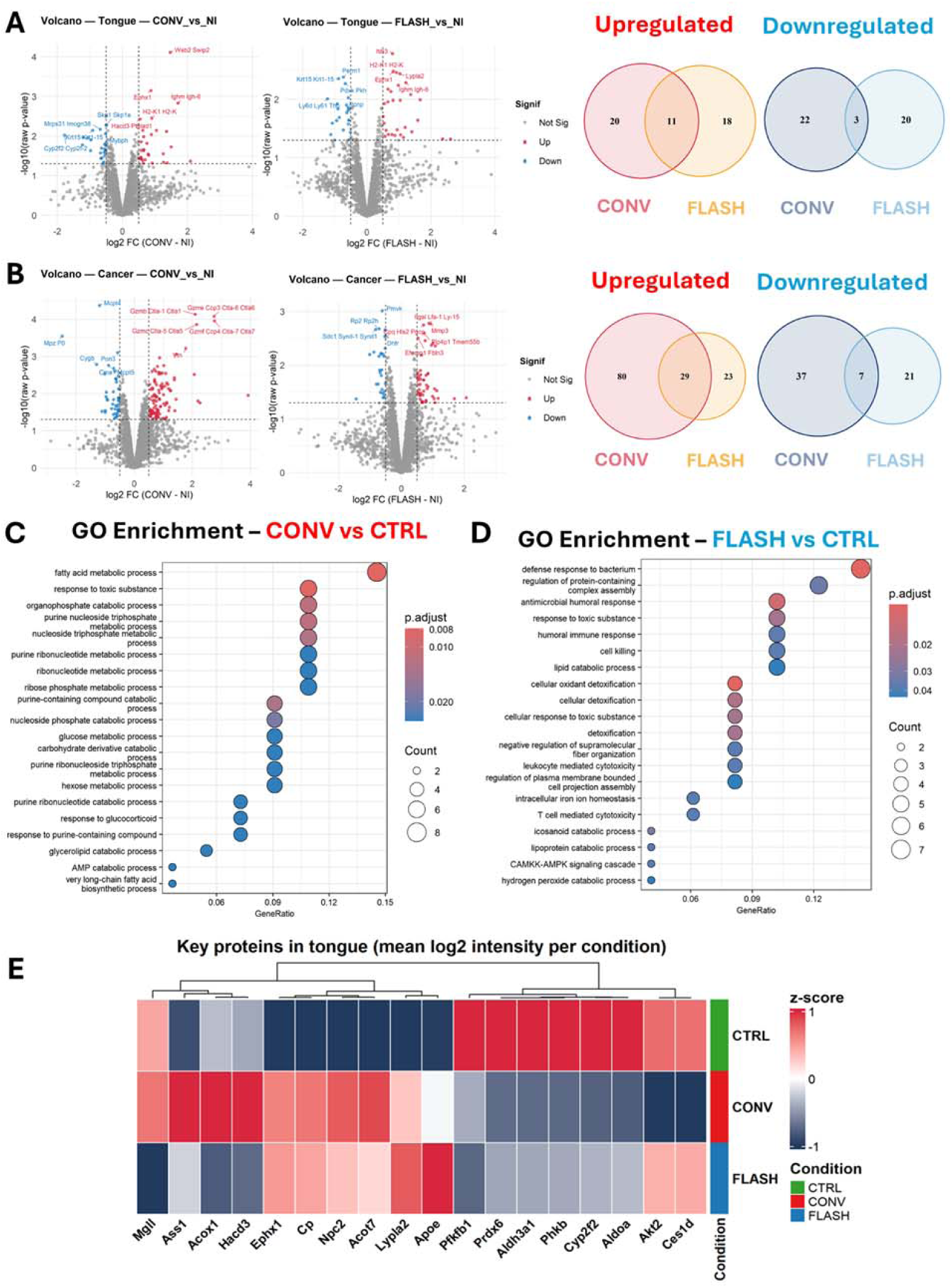
Distinct proteomic programs underlie CONV- and FLASH-specific responses in irradiated tongue tissue. (A) Volcano plots showing differential protein expression in tongue tissue comparing CONV vs CTRL and FLASH vs CTRL conditions. Significantly upregulated (red) and downregulated (blue) proteins are highlighted (adjusted *p*-value < 0.05). Right panels show Venn diagrams illustrating the overlap of upregulated and downregulated proteins between CONV and FLASH irradiation in tongue tissue. (B) Volcano plots of differential protein expression in contralateral MOC2 tumor tissue for CONV vs CTRL and FLASH vs CTRL comparisons. Corresponding Venn diagrams summarize shared and modality-specific up- and downregulated proteins. (C) Gene Ontology (GO) enrichment analysis of significantly regulated proteins in tongue tissue after CONV irradiation relative to CTRL. Enriched biological processes are predominantly associated with fatty acid metabolism, lipid catabolic pathways, and responses to toxic substances. (D) GO enrichment analysis of significantly regulated proteins in tongue tissue after FLASH irradiation relative to CTRL, revealing enrichment of immune and inflammatory responses, detoxification pathways, and cellular defense mechanisms, alongside more limited catabolic processes. (E) Heatmap of selected significantly regulated proteins associated with lipid metabolism, energy homeostasis, and oxidative stress in tongue tissue across CTRL, CONV, and FLASH conditions. Proteins are hierarchically clustered based on Z-scored abundance.

Notably, functional enrichment analysis revealed clear qualitative differences between CONV and FLASH-induced proteomic responses (Figure 4C–D). After CONV irradiation, significantly regulated proteins were predominantly enriched for pathways related to fatty acid metabolism and responses to toxic substances, consistent with the pronounced neutral lipid depletion and metabolic stress observed at the lipidomic level. After FLASH irradiation, lipid catabolic processes were also enriched, but the profile was dominated by immune defense pathways, including antimicrobial responses, cell killing, and T cell-mediated cytotoxicity, alongside cellular detoxification and oxidant defense mechanisms (Fig. 4D). Notably, icosanoid catabolic process and lipoprotein catabolic process were enriched exclusively in the FLASH condition, suggesting modality-specific engagement of lipid mediator and lipoprotein processing pathways.

Heatmap-based hierarchical clustering of selected significantly regulated proteins associated with lipid metabolism, energy homeostasis, and oxidative stress further resolved distinct and coherent response modules discriminating CONV and FLASH irradiation in tongue tissue (Figure 4E), revealing a catabolic and stress-driven program after CONV and a lipid-protective and remodeling-oriented program after FLASH. Enzymes involved in glycolysis and energy metabolism (Pfkfb1, Aldoa, Phkb) and antioxidant defense (Prdx6, Cyp2f2, Aldh3a1) were consistently downregulated after irradiation, with stronger suppression after CONV conditions, suggesting a more pronounced disruption of cellular energy production and redox capacity. CONV irradiation selectively suppressed a pro-survival kinase in the PI3K/Akt pathway (Akt2) and a triglyceride hydrolase essential for lipid droplet mobilization (Ces1d), linking the proteomic response directly to the triglyceride depletion observed at the lipidomic level. Acyl-CoA thioesterase 7 (Acot7) was induced after both modalities but more strongly after CONV, consistent with greater fatty acid flux from TG breakdown. Core detoxification functions, including iron transport (Cp) and epoxide hydrolysis (Ephx1), were comparably upregulated after both conditions. In contrast, FLASH irradiation preferentially induced a lipid transport protein involved in cholesterol and phospholipid redistribution during tissue repair (Apoe) and a lysophospholipase involved in membrane lipid homeostasis (Lypla2), consistent with active lipid remodeling. FLASH irradiation also suppressed monoacylglycerol lipase (Mgll), the terminal enzyme of glycerolipid hydrolysis (CONV vs FLASH, p = 0.025), further limiting TG catabolism relative to CONV. CONV irradiation also upregulated Npc2, a lysosomal cholesterol transport protein, suggesting mobilization of lipid droplet-derived cholesterol following TG depletion. Finally, a CONV-specific cluster comprising enzymes of amino acid metabolism (Ass1), peroxisomal fatty acid β-oxidation (Acox1), and fatty acid elongation (Hacd3) highlighted the catabolic and oxidative stress program driven by conventional irradiation. The same set of proteins analyzed in MOC2 tumor tissue showed no equivalent CONV versus FLASH polarization (Supplementary Figure 4B), indicating that the divergent metabolic programs observed here are specific to tongue tissue.

## Discussion

This study reveals that FLASH and CONV proton irradiation induce divergent lipid metabolic programs in normal tongue tissue, with the most pronounced differences localized to minor salivary glands, whereas tumors exhibited only modest lipidomic alterations across irradiation conditions. The differential impact of FLASH versus CONV irradiation on lipid profiles may originate in part from acute differences in lipid peroxidation. Prior studies have demonstrated that delivering radiation at ultra-high dose rates fundamentally alters the early physicochemical events of lipid oxidation. Notably, Froidevaux et al. showed in model membrane systems that FLASH irradiation fails to induce any detectable lipid peroxidation, whereas conventional dose-rate irradiation produces robust peroxidative damage^35^. This suppression appears selective for normal tissue contexts. In tumor cells, which often have high iron and reactive oxygen species (ROS) levels, FLASH irradiation can still trigger lipid peroxidation and ferroptotic cell death similarly to CONV irradiation. Vilaplana-Lopera et al. discussed that while FLASH did not significantly elevate lipid peroxidation in normal mouse tissues, it induced comparable levels of lipid peroxidation in cancer cells^42^. Consistent with this, Ni et al. demonstrated that FLASH radiation specifically abrogates lipid oxidase expression and oxidized low-density lipid generation in tumor macrophages, while standard radiation induces ROS-dependent lipid oxidation pathways^28^. Thus, the FLASH effect spares normal tissues by avoiding excessive lipid peroxidation yet maintains its efficacy in tumor tissues where cancer cell-intrinsic factors still drive lipid peroxidation and cell death. However, these acute peroxidative differences are short-lived. Portier et al. demonstrated that oxylipin differences between FLASH and CONV irradiation resolve within 24 hours, followed by continued remodeling at later timepoints^36^. Yet the functional consequences of FLASH irradiation persist well beyond this window. In the same head and neck cancer model used in the present study, Chowdhury et al. demonstrated that FLASH proton radiation ameliorates salivary gland damage and oral mucositis while maintaining tumor control^24^, with protection extending from reduced cell death and preserved aquaporin-5 expression at 14 days to decreased fibrosis at 28–60 days and improved salivary flow at 4–8 weeks. This temporal dissociation between the rapid resolution of acute lipid peroxidation and the sustained tissue protection suggests that biological mechanisms beyond the initial physicochemical events contribute to the FLASH effect. Our multi-omics analysis at two weeks post-irradiation, the same timepoint as the histopathological assessment of Chowdhury et al., offers a molecular window into this gap.

Our spatial lipidomics analyses using MALDI-MSI revealed that triglyceride signals in control tongue tissue were not uniformly distributed but were strongly enriched within discrete glandular compartments (Fig. 3A). Overlay with H&E-stained sections demonstrated that these TG species were highly concentrated in minor salivary glands, encompassing both serous and mucous acini, consistent with the lipid-rich secretory nature of these structures (Fig. 3A; Supplementary Fig. 3). Following irradiation, a marked and spatially restricted loss of TG signal was observed within these glandular regions. This depletion was particularly pronounced after CONV irradiation, where TG signals were strongly reduced or nearly absent in minor salivary glands. In contrast, FLASH irradiation was associated with partial preservation of TG signal intensity within the same acinar compartments. These data identify minor salivary glands as a major anatomical locus of radiation-induced triglyceride depletion and highlight their pronounced vulnerability to conventional irradiation, with partial protection conferred by FLASH. These spatial findings provide a lipid metabolic basis for the functional observations of Chowdhury et al. ^24^: the loss of neutral lipid reserves in glandular acini may directly compromise secretory capacity, and the preservation of TG stores after FLASH is consistent with the maintained salivary function they reported. Integrating these spatial lipidomic findings with our proteomic data reveals a coherent mechanistic picture of the TG depletion cascade.

The dramatic depletion of triglycerides in CONV-irradiated tissue was accompanied by the loss of carboxylesterase 1d (Ces1d), a triglyceride hydrolase that mobilizes fatty acids from lipid droplets^43^(Fig. 4E). The fatty acids released by this mobilization were then catabolized through peroxisomal β-oxidation, as indicated by the striking upregulation of acyl-CoA oxidase 1 (Acox1), the rate-limiting enzyme of this pathway. Acox1-driven β-oxidation is known to generate toxic levels of ROS^44^, linking TG mobilization directly to oxidative damage. The downregulation of aldehyde dehydrogenase 3A1 (Aldh3a1), which detoxifies lipid peroxidation-derived aldehydes such as 4-hydroxynonenal, was observed after both modalities but more pronounced after CONV, consistent with a greater lipid peroxidation burden under conventional irradiation as reported by Froidevaux et al.^35^. Acyl-CoA thioesterase 7 (Acot7), which hydrolyzes long-chain acyl-CoAs to free fatty acids^45^, was also more strongly induced after CONV than FLASH, consistent with TG breakdown. This interpretation is further supported by gene ontology analysis, which revealed enrichment of fatty acid metabolic process and glycerolipid catabolism specifically after CONV irradiation (Fig. 4C).

Conversely, FLASH irradiation suppressed monoacylglycerol lipase (Mgll) ^46^, the terminal enzyme of glycerolipid hydrolysis, relative to CONV, suggesting that FLASH actively limits the complete degradation of TG stores. In parallel, FLASH irradiation preferentially induced apolipoprotein E (Apoe), a lipid transport protein that mediates the redistribution of triglycerides and phospholipids via VLDL and HDL particles (Fig. 4E). Apoe plays a well-established role in mobilizing lipid reserves to sites of tissue repair^47^, and its upregulation after FLASH is consistent with active redistribution rather than catabolism of TG stores. The Apoe-mediated lipid redistribution described above could provide a mechanistic link between TG preservation and the second major finding of this study: active membrane phospholipid remodeling after FLASH irradiation. The increase in membrane phospholipids observed in the FLASH group, most prominently in phosphatidylethanolamine (PE: 28.0% vs 10.9% of species upregulated), a phospholipid enriched in mitochondrial membranes and intracellular organelles, and to a lesser extent phosphatidylcholine (PC: 19.1% vs 14.2%), extends beyond passive damage avoidance predicted by the oxygen depletion hypothesis and the radical recombination hypothesis^48–50^. We interpret this phospholipid surge as evidence that FLASH triggers active membrane repair and reconstruction programs. FLASH-exposed cells may be proactively synthesizing and incorporating new phospholipids to repair sub-lethal membrane injuries. FLASH irradiation also upregulated a broader spectrum of lysophosphatidylcholine (LPC) and lysophosphatidylethanolamine (LPE) species compared to CONV (Fig. 2C). The concurrent elevation of both intact membrane phospholipids and their lysophospholipid derivatives (LPC, LPE) specifically after FLASH irradiation is consistent with active removal of damaged acyl chains from membrane phospholipids and subsequent de novo synthesis of replacement PC and PE. Notably, Lands cycle reacylation reshuffles acyl chains within existing phospholipids without increasing total pool size; the net gains in PC and PE abundance observed at two weeks post-irradiation therefore more likely reflect de novo synthesis during the tissue regeneration phase. Lysophospholipase 2 (Lypla2), a cytosolic enzyme that hydrolyzes lysophospholipids^51^, was upregulated after FLASH (Fig. 4E). The upregulation of Lypla2 may serve to limit the pro-inflammatory accumulation of lysophospholipids generated by damaged acyl chain removal, rather than participating directly in membrane remodeling itself^52^. Consistent with this interpretation, unlike the glycerolipid catabolism enriched after CONV, FLASH-specific enrichment of lipoprotein catabolic process and icosanoid catabolic process suggests that lipid breakdown after FLASH serves a remodeling and resolution function rather than an emergency energy supply.

Beyond lipid-specific enzymes, the proteomic data revealed additional markers of the divergent tissue responses. Argininosuccinate synthase 1 (Ass1), which synthesizes arginine to support high-output nitric oxide production by iNOS^53^, was upregulated uniquely after CONV irradiation, consistent with the enrichment of response to toxic substance pathways observed after CONV (Fig. 4C). This suggests that conventional irradiation created an inflammatory milieu in which arginine–NO pathways are overactive. Akt2, a kinase central to pro-survival signaling, was significantly down-regulated only after CONV irradiation^54^, indicating a collapse of pro-survival signaling. Not all proteomic changes were modality-specific. Ceruloplasmin (Cp) and epoxide hydrolase 1 (Ephx1) were comparably upregulated after both CONV and FLASH (Fig. 4E), indicating that both modalities engage a common detoxification response to oxidative stress. This suggested that the divergence between CONV and FLASH thus lies not just in the absence of a damage response after FLASH, but in the additional engagement of lipid-protective and remodeling programs. In tumor tissue, no such polarized proteomic response was observed; both modalities activated similar anti-tumor and cell-killing pathways (Supplementary Fig. 4A), consistent with preclinical and emerging clinical evidence that FLASH irradiation maintains therapeutic efficacy against tumors^5,24–27,55^.

In summary, our findings suggest that the FLASH effect extends beyond a reduction in initial physicochemical damage to include active reprogramming of lipid metabolism in irradiated normal tissues. We term this the Lipid Metabolic Reprogramming Hypothesis. This hypothesis complements existing physicochemical models of the FLASH effect by adding an active metabolic dimension to the predominantly acute mechanisms proposed to date. This reprogramming was most pronounced in the highly radiosensitive salivary glands, suggesting that FLASH may confer selective protection commensurate with the inherent vulnerability of tissues.

In conclusion, this integrated lipidomic, spatial lipidomic, and proteomic analysis reveals molecular signatures that distinguish FLASH from conventional proton irradiation and identifies lipid metabolic reprogramming as a putative mechanism of the FLASH effect. The selective vulnerability of radiosensitive glandular structures to lipid metabolic disruption, and their preferential protection under FLASH conditions, may inform strategies to mitigate normal tissue toxicity in radiotherapy for head and neck cancer. Functional validation of this metabolic reprogramming, characterization of its temporal kinetics, and extension to fractionated regimens will be essential next steps toward clinical translation.

## Author Contributions

S.A.: Conceptualization, Methodology, Formal analysis, Visualization, Writing (original draft, review and editing). M.D.: Formal analysis, Writing (review and editing). M.Se. (Sertorio): Methodology (animal model, irradiation), Resources, Writing (review and editing). Md.M.R.: Methodology. M.S.A.: Methodology. C.Z.: Methodology, Writing (review and editing). L.T.: Formal analysis. K.W.: Writing (review and editing). K.K.: Supervision. T.K.: Supervision. G.B.: Project administration. M.V.: Project administration. R.A.S.: Conceptualization, Supervision, Funding acquisition, Writing (review and editing). K.N.: Supervision. M.Se. (Setou): Conceptualization, Supervision, Funding acquisition, Resources, Writing (review and editing).

## Data Availability

The lipidomics, proteomics, and MALDI-MSI datasets generated during this study are available from the corresponding author upon reasonable request. Data processing scripts (R v.4.2) used for lipidomic and proteomic analyses are available from the corresponding author upon reasonable request.

