## Supplementary Table 1, Supplementary Figures 1-4 for "Reprogramming of Lipid Metabolism by FLASH Radiotherapy Selectively Protects Radiosensitive Normal Tissues"

**Supplementary Table 1. Proton irradiation parameters for FLASH and conventional dose-rate delivery.**

| **Parameter** | **FLASH** | **Conventional** |
| --- | --- | --- |
| ***Beam delivery*** | | |
| Particle | Proton | Proton |
| System | ProBeam PBS gantry (Varian) | ProBeam PBS gantry (Varian) |
| Beam energy (MeV) | 250 | 244 |
| Delivery mode | Single-layer transmission | Single-layer transmission |
| Beam continuity | Continuous (no inter-spot pause) | Continuous (no inter-spot pause) |
| ***Dose specification*** | | |
| Prescribed dose (Gy) | 10 | 10 |
| Dose rate (Gy/s) | 100 | 1 |
| Dose tolerance (%) | ±3 | ±3 |
| Dose-rate tolerance (%) | ±5 | ±5 |
| ***Field geometry*** | | |
| Field size, 95% isodose (mm²) | 25 × 23 | 25 × 23 |
| Number of spots | 30 | 30 |
| Spot size σ (mm) | 4.0 | 4.0 |
| Uniformity tolerance (%) | ±5 | ±5 |
| Flatness / symmetry tolerance (%) | ≤5 / ≤5 | ≤5 / ≤5 |
| Gantry angle (°) | 0 | 0 |
| ***Dosimetry*** | | |
| Dose formalism | IAEA TRS-398 | IAEA TRS-398 |
| Ion chamber | Advanced Markus (PTW) | Advanced Markus (PTW) |
| Electrometer | IBA Dose1 | IBA Dose1 |
| Radiochromic film | Gafchromic EBT3 (Ashland) | Gafchromic EBT3 (Ashland) |
| Ion recombination correction (%) | <1.0 | <1.0 |
| Calorimeter agreement (%) | ±1.0 (graphite) | ±1.0 (graphite) |
| Measurement depth, WED (cm) | 1.0 | 1.0 |
| Online verification | Distal ion chamber | Distal ion chamber |
| ***Alignment*** | | |
| Primary method | Gantry laser (±1 mm) | Gantry laser (±1 mm) |
| Field verification | Radiochromic film in jig | Radiochromic film in jig |
| Image guidance | Orthogonal kV (subset) | Orthogonal kV (subset) |

All irradiations were performed using the ProBeam Pencil Beam Scanning (PBS) Gantry (Varian Medical Systems, Palo Alto, CA, USA). A single fraction of 10 Gy was delivered to each mouse. Dose rate was determined as the ratio of total dose to total field delivery time from machine log files. Abbreviations: PBS, Pencil Beam Scanning; WED, water-equivalent depth; kV, kilovoltage; IAEA, International Atomic Energy Agency; TRS, Technical Report Series.

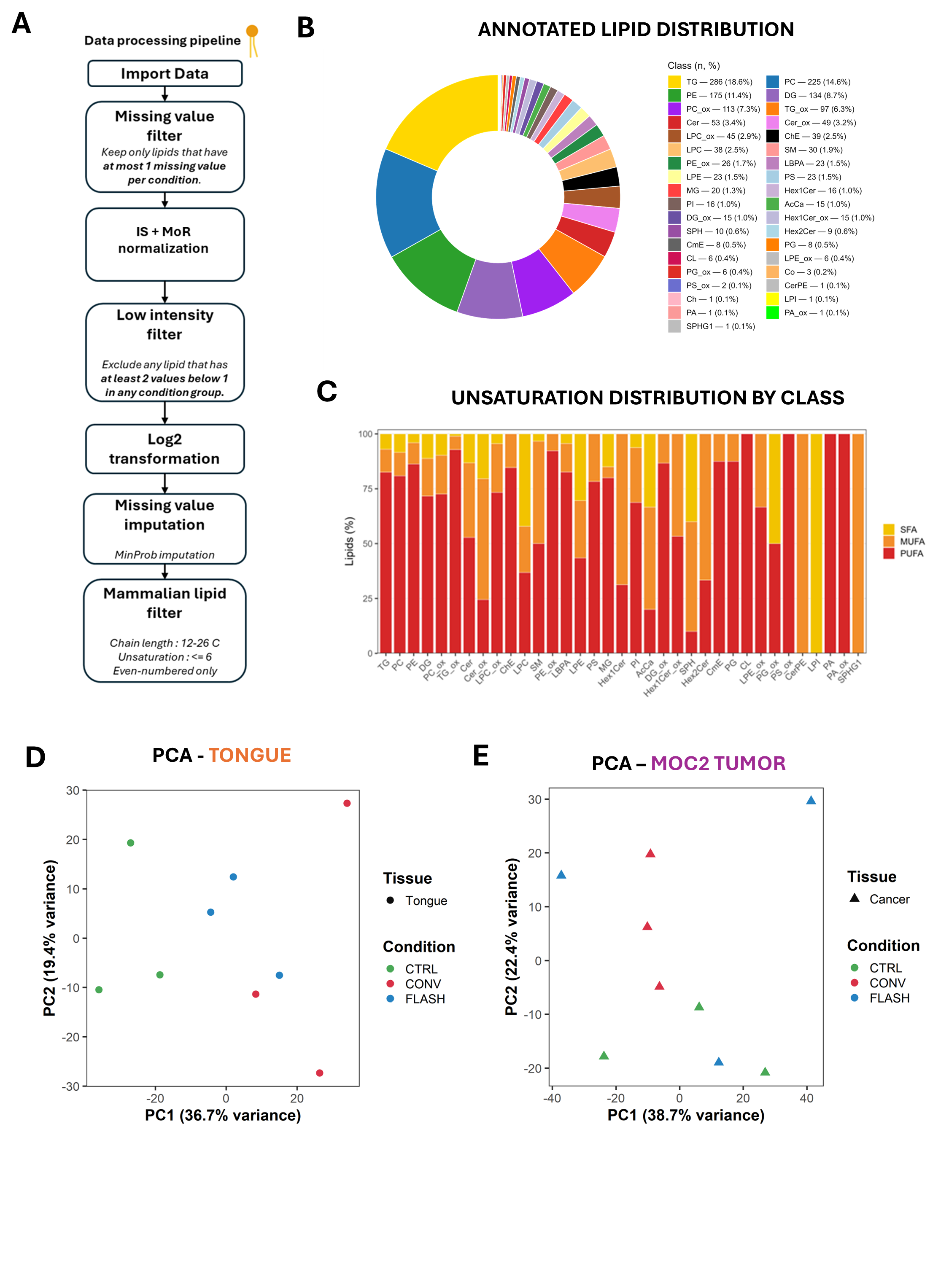

**Supplementary Figure 1. Overview of the lipidomic data processing pipeline and baseline lipidome characteristics.** (A) Schematic representation of the lipidomic data processing workflow (B) Global annotated lipid class distribution across all samples, expressed as the number and percentage of detected lipid species per class. (C) Distribution of fatty acyl chain unsaturation levels within each lipid class, showing the relative contributions of saturated (SFA), monounsaturated (MUFA), and polyunsaturated (PUFA) species and illustrating class-specific biochemical properties. (D) Principal component analysis (PCA) of lipidomic profiles in tongue tissue, showing separation of samples according to irradiation condition (CTRL, CONV, FLASH). (E) PCA of lipidomic profiles in MOC2 tumor tissue, revealing more limited separation between experimental conditions compared with tongue tissue.

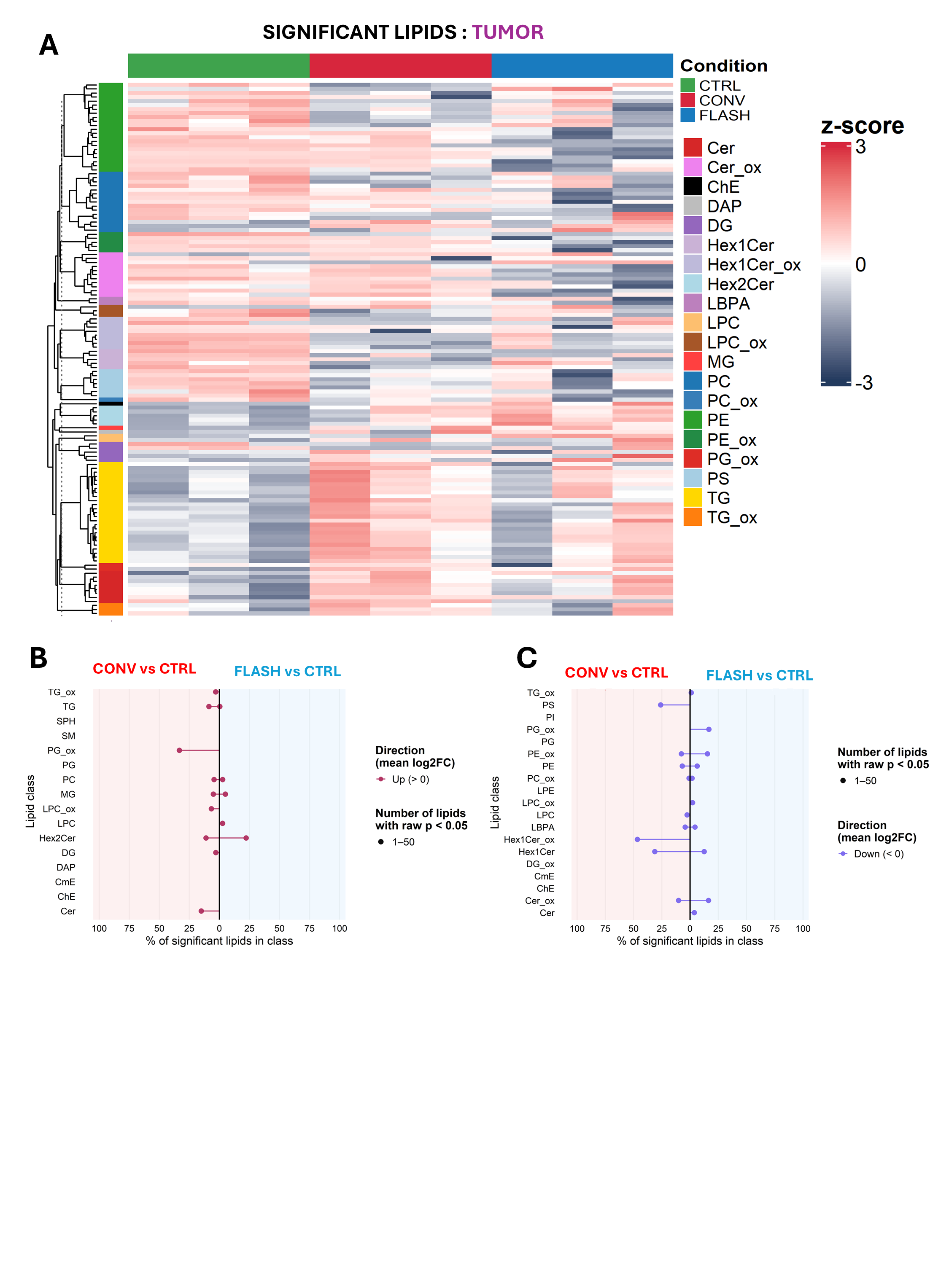

**Supplementary Figure 2. Attenuated and heterogeneous lipidomic remodeling in MOC2 tumor tissue following irradiation.** (A) Lipid class–based hierarchical clustering heatmap of significantly regulated lipid species in MOC2 tumor tissue across CTRL, CONV, and FLASH conditions. Lipid abundances are shown as row-wise z-scores, and lipid species are annotated by class (color-coded sidebar).(B) Lipid class–level analysis of significantly *upregulated* lipid species in tumors. Butterfly plots depict the percentage of significantly upregulated lipids within each class for CONV vs CTRL (left) and FLASH vs CTRL (right) comparisons. Point size reflects the number of significant species per class (raw p < 0.05), and direction is defined by the mean log₂ fold change. (C) Lipid class–level analysis of significantly *downregulated* lipid species in tumors, displayed as butterfly plots for CONV vs CTRL and FLASH vs CTRL comparisons, using the same representation as in panel B.

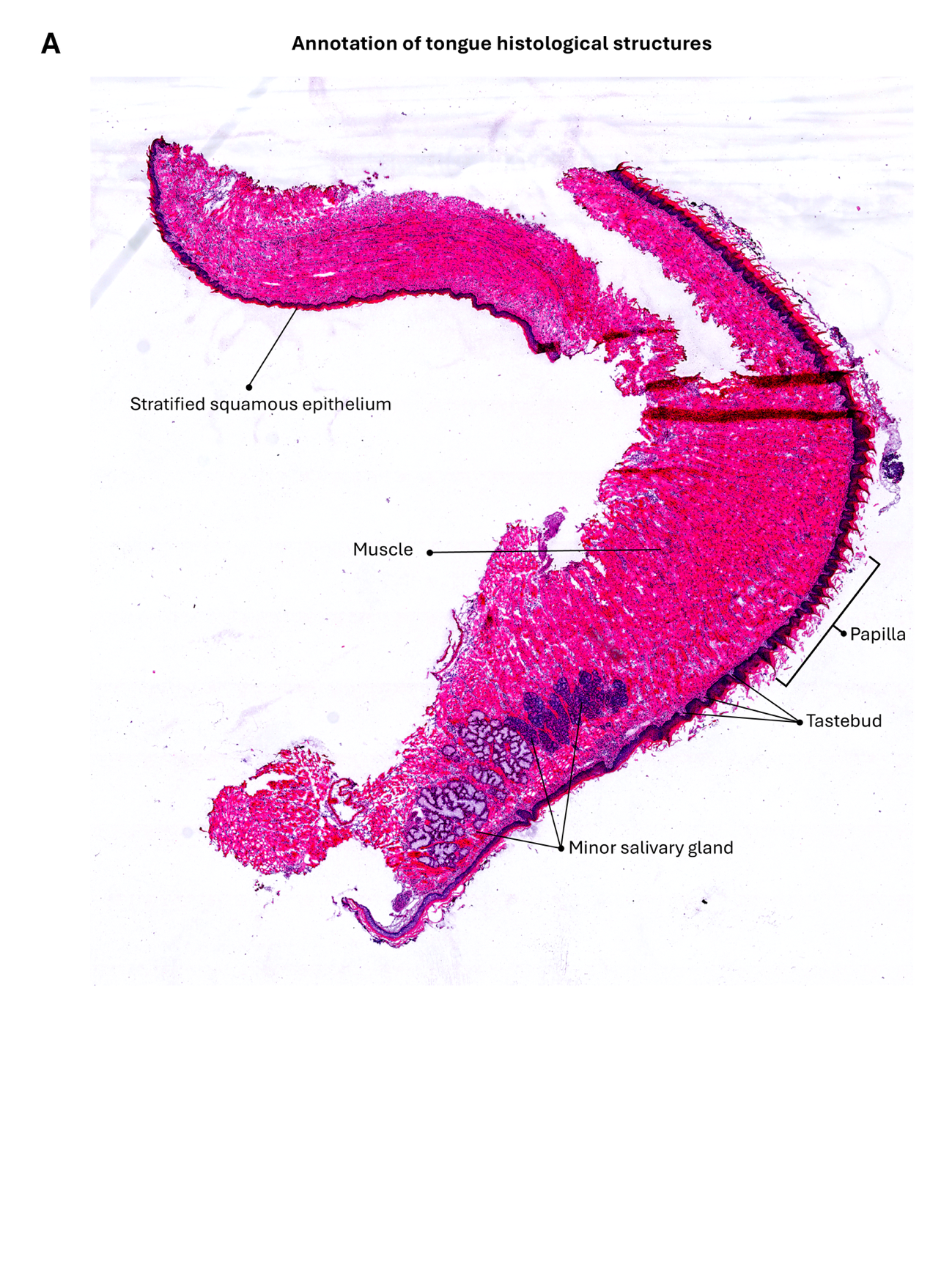

**Supplementary Figure 3. Histological annotation of mouse tongue tissue.** Representative H&E-stained section of mouse tongue with annotation of major histological structures. Key anatomical features are indicated: stratified squamous epithelium (surface layer), underlying skeletal muscle fibers, lingual papillae (dorsal surface projections), taste buds (sensory structures within papillae), and minor salivary glands (serous and mucous acini). This reference image provides anatomical context for interpretation of MALDI-MSI data presented in Figure 3.

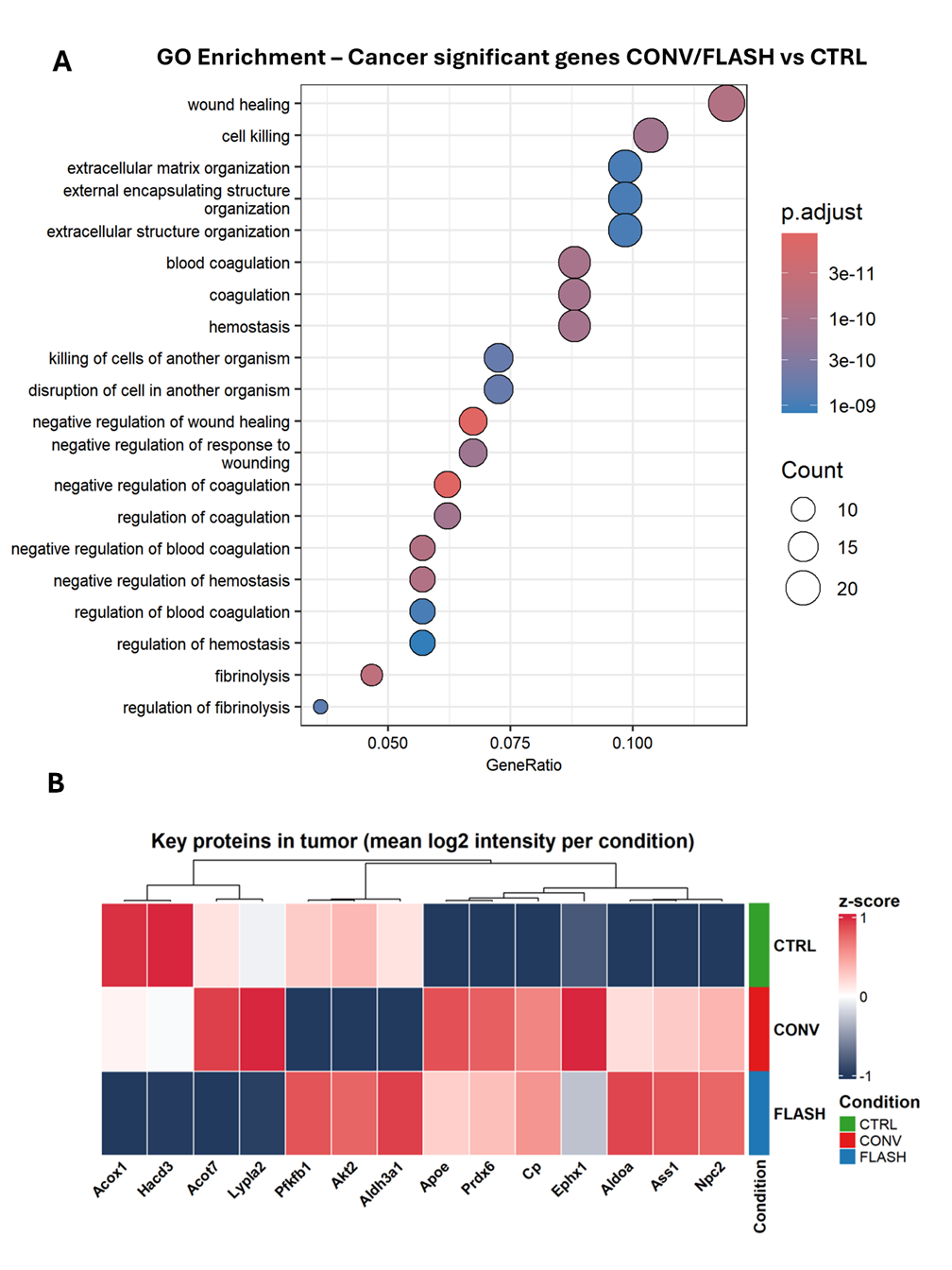

**Supplementary Figure 4. Proteomic responses in MOC2 tumor tissue following CONV and FLASH irradiation.** (A) Dot plot of Gene Ontology (biological process) terms enriched among proteins significantly regulated in tumor tissue after CONV and FLASH irradiation compared with CTRL. Enrichment highlights pathways related to wound healing, extracellular matrix organization, coagulation/hemostasis, and cell killing. Dot size represents the number of proteins contributing to each term (Count), color indicates adjusted p-value (p.adjust), and the x-axis shows the GeneRatio. (B) Heatmap of selected proteins associated with lipid metabolism, energy homeostasis, and oxidative stress in MOC2 tumor tissue across CTRL, CONV, and FLASH conditions. Values are displayed as row-wise z-scores of mean log₂ intensity per condition; hierarchical clustering was performed on columns. This panel provides a tumor counterpart to the tongue-tissue heatmap in Figure 4E.
